# PRS-Net: Interpretable polygenic risk scores via geometric learning

**DOI:** 10.1101/2024.02.13.580211

**Authors:** Han Li, Jianyang Zeng, Michael P. Snyder, Sai Zhang

## Abstract

Polygenic risk score (PRS) serves as a valuable tool for predicting the genetic risk of complex human diseases for individuals, playing a pivotal role in advancing precision medicine. Traditional PRS methods, predominantly following a linear structure, often fall short in capturing the intricate relationships between genotype and phenotype. We present PRS-Net, an interpretable deep learning-based framework designed to effectively model the nonlinearity of biological systems for enhanced disease prediction and biological discovery. PRS-Net begins by deconvoluting the genomewide PRS at the single-gene resolution, and then it encapsulates gene-gene interactions for genetic risk prediction leveraging a graph neural network, thereby enabling the characterization of biological nonlinearity underlying complex diseases. An attentive readout module is specifically introduced into the framework to facilitate model interpretation and biological discovery. Through extensive tests across multiple complex diseases, PRS-Net consistently outperforms baseline PRS methods, showcasing its superior performance on disease prediction. Moreover, the interpretability of PRS-Net has been demonstrated by the identification of genes and gene-gene interactions that significantly influence the risk of Alzheimer’s disease and multiple sclerosis. In summary, PRS-Net provides a potent tool for parallel genetic risk prediction and biological discovery for complex diseases.

## 1 Introduction

Complex human diseases display polygenicity in their genetic architectures, characterized by a multitude of common genetic variants with minor individual effects accumulatively influencing the disease risk^1–4^. The polygenic risk scores (PRSs) are developed to quantitatively characterize the genetic susceptibility of individuals to specific traits or complex diseases based on the common genetic variants^5–7^. This methodology empowers the early deployment of targeted therapeutic interventions and facilitates the practice of personalized medicine^8–10^.

PRS is typically calculated using the summary statistics derived from genome-wide association studies (GWAS)^11–17^, a widely-used statistical method for identifying disease-associated genetic variants^18–20^. While GWAS can identify disease risk genetic variants, such as single nucleotide polymorphisms (SNPs), that exhibit significant differences in frequencies between cases and controls, these variants tend to have modest individual effects on the phenotype, resulting in limited prediction capability. In an effort to enhance predictive modeling, various statistical methods have been applied to aggregate the effects of individual SNPs. The widely adopted method for calculating PRS, exemplified by PLINK^21^ and PRSice^12^, is known as clumping and thresholding (C+T)^11^, which involves summing allele counts weighted by effect sizes estimated from GWAS. More recent approaches like LDpred2^16^ utilize Bayesian modeling to infer the posterior mean effect size of each marker by incorporating prior information on effect sizes and linkage disequilibrium (LD) data from an external reference panel. Similarly, lassosum2^17^ estimates PRS using summary statistics and a reference panel within a penalized regression framework. With the notable increase in dataset sample sizes for GWAS, these methods have achieved enhanced predictive power^22^. Nonetheless, these techniques primarily rely on univariate effect sizes derived from linear GWAS models, thus often overlook potential non-linear associations between genetic factors and phenotypes, which can undermine their predictive performance.

Efforts have also been made to construct models capable of capturing non-linear interactions in PRS calculation. These include tree-based methods like random forests^23, 24^, gradient boosting^25, 26^, and AdaBoost^27, 28^, as well as deep learning-based techniques such as multiple-layer perceptrons (MLP)^29^ and convolutional neural networks^30^. However, these methods only take a limited number of variants as their input, and lack the integration of versatile prior biological knowledge. Indeed, these approaches have demonstrated either comparable or, in many cases, less effective performance in predicting phenotypes when compared to linear models^31, 32^.

In this study, we propose PRS-Net, a geometric deep learning-based approach designed to effectively model the intricate non-linear relationships among genetic factors such as genes in predicting the disease risk, thus delivering more accurate and robust PRSs. Based on the summary statistics of GWAS, PRS-Net first maps PRS onto a gene-gene interaction (GGI) network through the derivation of gene-level PRSs using the C+T method. Subsequently, a graph neural network is employed to iteratively update the embedding of the genes via performing message passing on the GGI network, thus capturing the complex GGIs from the network. An attentive readout module is then introduced to provide interpretable PRS predictions. PRS-Net also integrates a mixture-of-expert module^33^ designed to enhance the accuracy of PRS predictions when dealing with multi-ancestry datasets. Our comprehensive evaluation encompasses six complex diseases extracted from the UK Biobank database^34^, including Alzheimer’s disease, atrial fibrillation, rheumatoid arthritis, multiple sclerosis, ulcerative colitis, and asthma. The results consistently demonstrated the superiority of PRS-Net over baseline methods, including PLINK^21^, PRSice2^14^, LDpred-2^16^, and lassosum2^17^ in PRS prediction. Notably, through case studies focused on Alzheimer’s disease and multiple sclerosis, we illustrated that PRS-Net provided biologically meaningful interpretability by identifying specific genes and GGIs that significantly influence disease risk. In summary, PRS-Net stands as a potent and innovative tool for precise PRS prediction, addressing the limitations of current linear models and offering a more comprehensive approach to unraveling the genetic underpinnings of complex traits and diseases.

## 2 Method

In this section, we present our proposed framework for PRS estimation (Fig. 1), covering the establishment of the GGI network, the derivation of gene-level PRS, and the architecture of PRSNet.

**Fig. 1:**
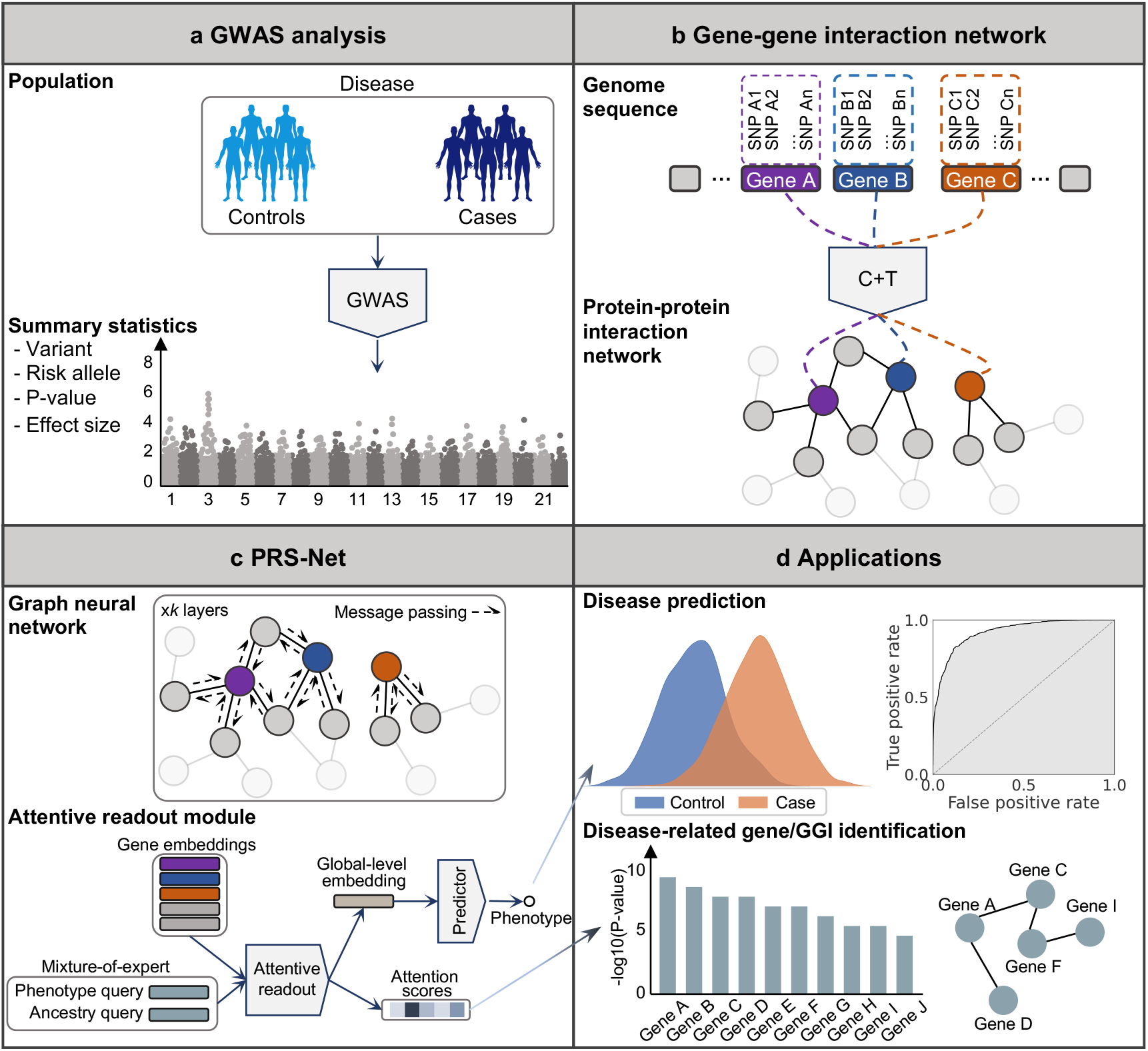
An illustrative diagram of PRS-Net. **a** The proposed framework is based on summary statistics, including variants, risk alleles, P-values, and effect sizes derived from GWAS. **b** A gene-gene interaction network is constructed based on the protein-protein interaction network. Gene-level PRSs are calculated with the C+T method to serve as the node features for the nodes within the network. **c** A graph neural network is employed to update node features via message passing and subsequently an attentive readout module is applied to provide interpretable PRS predictions. **d** The PRS-Net can be applied for disease prediction and disease-related gene/GGI identification.

### 2.1 GGI network

It is widely recognized that the disease phenotype is not solely determined by individual genes but rather involves the intricate interactions among multiple genes, which can exhibit additive or non-additive genetic relationships^35–37^. Additive genetic interactions manifest when the cumulative effects of genes jointly contribute to a specific phenotype. Furthermore, there are increasing studies highlighting the significance of non-additive genetic interactions^38–40^. Epistasis is a prominent example of non-additive genetic interaction, which occurs when the impact of a gene mutation depends on the presence or absence of mutations in one or more other genes^41–43^. We establish a GGI network that empowers PRS-Net to capture the intricate genetic relationships that are potentially associated with the target phenotypes (Fig. 1b).

We construct our GGI network based on the protein-protein interactions derived from the STRING database^44^, as protein-protein interactions represent potent indicators of functional relationships between genes. Formally, we construct a GGI network, denoted as ***𝒢*** = (***𝒱***,***ℰ***), where stands ***𝒱*** for the set of nodes and ***ℰ*** stands for the set of edges. Each node *v*_*i*_*∈* ***𝒱*** stands for a coding gene and each edge (*v*_*i*_, *v*_*j*_) *∈* ***ℰ*** stands for an interaction between nodes *v*_*i*_ and *v*_*j*_ derived from the STRING database^44^. Note that, we add a self-loop (*v*_*i*_, *v*_*i*_) for each node *v*_*i*_ *∈* ***𝒱***. This network construction results in a GGI network encompassing 19,836 coding genes and 250,236 interactions.

Upon deriving the GGI network, we proceed to compute gene-level PRSs for the genes within the network using a C+T approach^11, 21^. More precisely, for each gene in the network, we focus on the SNPs falling within a designated range, spanning from its transcription start site *L* to its transcription end site + *L*. In our tests, we set *L* to 10 kilobases (KB), thereby encompassing the SNPs situated in non-coding regions, such as the promoters of the genes. Subsequently, for each gene, we perform LD clumping on the associated SNPs from the GWAS data, utilizing the LD information estimated in the target data. Following this, we filter the SNPs based on a specific P-value threshold. The gene-level PRSs are then derived by multiplying the genotype matrix by the effect sizes obtained from the GWAS data, and then dividing this by the number of allele observations for each gene. For the LD clumping process, we set the LD threshold *R*^2^ to 0.5 and the physical distance threshold to 250 KB. As for the thresholding step, we set the P-values to 1*e*^*−*5^, 1*e*^*−*4^, 1*e*^*−*3^, 1*e*^*−*2^, 5*e*^*−*2^, 0.1, 0.2, 0.3, 0.5, and 1, respectively. This process results in the computation of eleven PRSs for each gene, which serves as their initial features. We denote the initial feature of *v*_*i*_ *∈* ***𝒱*** as ***h***_*i*_ *∈* ***H***, where ***H*** *∈* ℝ^|***𝒱***|*×*11^ and |***𝒱***| stands for the number of genes in ***𝒢***.

### 2.2 PRS-Net

#### Graph neural network

We harness the power of a graph neural network to capture the complex interactions among genes within our established GGI network (Fig. 1c). In our tests, we specifically opt for a graph isomorphism network (GIN)^45^ due to its proven theoretical and experimental expressiveness. Formally, we first encode the initial feature of nodes, denoted as ***H***, by employing an MLP in the following manner:

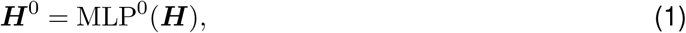

where ***H***^0^ *∈*ℝ^|***𝒱***|*×D*^ and D is the dimension of hidden states. Subsequently, we apply multiple GIN layers to iteratively update the representation of each node by aggregating the representations of

its neighbors, as depicted below:

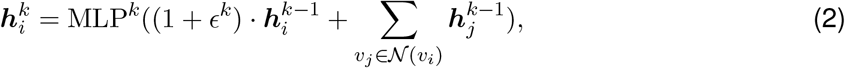

where 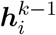 is the hidden states of *v*_*i*_ at the (*k−*1)-th layer, *𝒩* (*v*_*i*_) stands for the neighbors of *v*_*i*_ in the GGI network, 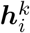 stands for the updated hidden states of *v*_*i*_ at the *k*-th layer, MLP^*k*^ is the MLP at the *k*-th layer, and *ϵ* stands for a learnable variable. Following *k* iterations of aggregation, each gene effectively encapsulates the interaction information within its *k*-hop neighborhood.

#### Attentive readout module

To make predictions for each data sample, we derive the global-level representation for each sample through an attentive readout module, illustrated as follows:

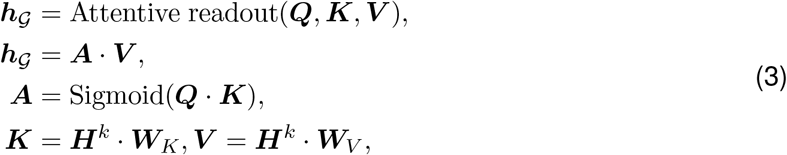

where ***W***_*K*_ *∈* ℝ^*D×D*^ and ***W***_*V*_ *∈* ℝ^*D×D*^ stand for trainable projection matrices to derive the key (i.e., ***K***) and value (i.e., ***𝒱*** ) matrices, respectively. ***Q*** *∈* ℝ^1*×D*^ stands for a trainable query vector. Sigmoid stands for the sigmoid function. ***A*** *∈* ℝ^1*×*|***𝒱***|^ stands for the attention scores, with elevated scores signifying a greater significance of the associated genes. ***h***_*𝒢*_*∈* ℝ^1*×D*^ stands for the global-level representation.

After deriving the global-level representation, we employ an MLP to derive the final prediction, denoted as 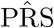, as follows:

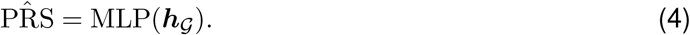

Additionally, we implement a mixture-of-expert module^33^ to effectively handle datasets that encompass data samples from multiple ancestries. More specifically, we introduce a specialized attentive readout module for each distinct ancestry. These dedicated attentive readout modules are activated when processing data from individuals with specific ancestral origins. To illustrate, when dealing with input samples of Western European ancestry, we derive the ancestry-specific global-level representation as follows:

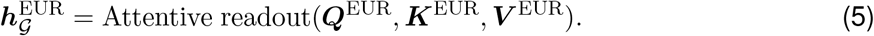

The ancestry-specific readout module is designed to capture the unique knowledge pertaining to each ancestry in relation to the disease. In addition, we introduce another shared readout module to capture disease-related knowledge that holds general applicability across all ancestries:

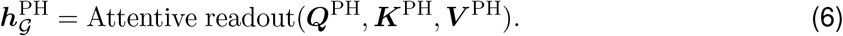

Then, we derive the final global-level representation by combining the aforementioned two representations:

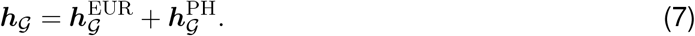

The process for deriving global-level representations of individuals from other ancestries follows a similar approach. The final PRS prediction can be computed with equation 4, utilizing the derived global-level representation. We refer to the single-ancestry variation as PRS-Net and the multiple-ancestry variation as PRS-Net_MA_.

## 3. Results

### 3.1 PRS-Net outperforms baseline methods in PRS prediction

We extracted genotype-phenotype data from the UK Biobank database^34^ for six different complex diseases, which encompassed Alzheimer’s disease, atrial fibrillation, rheumatoid arthritis, multiple sclerosis, ulcerative colitis, and asthma. ICD-10 codes^46^ were employed to define the disease phenotypes (Supplementary Table 1). For our primary experiments, we focused exclusively on individuals of Western European ancestry due to the insufficient size of the non-European ancestry population, which did not provide an adequate amount of training data (Supplementary Table 2). Following a quality control process, each disease dataset consisted of roughly 411,000 individuals (Supplementary Note 1.1). To prevent data leakage, we ensured that none of the GWAS were conducted on samples from the UK Biobank database (see Data availability). For each disease dataset, we randomly partitioned it into training, validation, and test sets with a ratio of 8:1:1. To evaluate the performance of PRS-Net, we compared it against several previously proposed methods, such as C+T-based methods (PLINK^21^ and PRSice2^14^), lassosum2^17^, and three variations of LDpred2^16^ (LDpred2-auto, LDpred2-grid, and LDpred2-inf), utilizing the area under the receiver operating characteristic curve (AUROC) as the metric. To ensure a rigorous and equitable comparison, we utilized LD matrices estimated from European populations within the 1000 Genomes Project^47^ across all methods in our study. Our results were based on three independent runs with different random seeds to ensure robustness and reliability. The results revealed that PRS-Net consistently outperformed all baseline methods on all disease datasets, resulting in relative improvements ranging from 0.5% to 3.7%. Interestingly, the largest improvements were obtained for two autoimmune diseases, i.e., ulcerative colitis (with a relative improvement of 3.0%) and multiple sclerosis (with a relative improvement of 3.7%), reinforcing the observed nonadditivity of genomic factors underlying these diseases^38, 48–50^. Altogether, our data demonstrates that PRS-Net possesses the capacity to capture more intricate associations between genotypes and phenotypes that are beyond the reach of previously proposed linear models.

We utilized the Aalen-Johansen estimator^51^ to estimate the disease occurrence over a lifetime for individuals categorized into high-risk and low-risk groups, as determined by the PRSs estimated by PRS-Net and baseline methods. High-risk individuals were defined as those with the highest 5% of PRSs, while low-risk individuals were identified as those with the lowest 5% of PRSs. The cumulative incidence plots revealed that individuals classified as high-risk by PRS-Net generally exhibited a heightened risk of disease throughout their lifetime compared to baseline methods, especially for ulcerative colitis, asthma, rheumatoid arthritis, and multiple sclerosis (Fig. 3a). Conversely, those categorized as low-risk by PRS-Net tended to maintain a lower risk of all diseases over their lifetime in comparison to baseline methods (Supplementary Fig. 1). These findings underscore the potential of PRS-Net as a powerful tool for individual risk stratification.

Next, we assessed the performance of PRS-Net and our multiple-ancestry model, PRS-Net_MA_, on a dataset comprising individuals from diverse ancestral backgrounds. Specifically, we curated a mixed-ancestry dataset encompassing Western European, South Asian, and African for asthma, which provides a reasonable number of asthma cases (over 1,000) for each ancestry (Supplementary Table 2). The results revealed that PRS-Net outperformed baseline methods on the mixed ancestry and South Asian ancestry test sets, indicating that the PRS-Net trained solely on the Western European ancestry dataset captured the underlying disease biology independent of different ancestries (Fig. 3b). Additionally, PRS-Net_MA_ demonstrated superior performance when compared to PRS-Net on the mixed ancestry, Western European ancestry, and African ancestry test sets (Fig. 3b). These findings underscored the ability of PRS-Net_MA_ to leverage the multiancestry dataset effectively, enhancing its portability in estimating PRS for individuals from diverse ancestral backgrounds.

### 3.2 PRS-Net identifies disease-related genes and GGIs for Alzheimer’s disease and multiple sclerosis

Following the demonstration of the superior performance of PRS-Net in predicting PRS, we sought to explore its capability to identify risk genes and GGIs underlying complex diseases. Alzheimer’s disease, a progressively degenerative condition, has been the subject of extensive research for many years, leading to the identification of numerous genes associated with the disease^52–56^. We employed PRS-Net to identify disease-related genes and GGIs, with the expectation that our findings would align with prior research outcomes. Specifically, we first applied the Mann–Whitney U test^57^ to each gene within our constructed GGI network, assessing whether the attention scores associated with the gene for individuals with Alzheimer’s disease were notably higher than those of the control group. This analysis yielded a gene set comprising 309 genes with compelling statistical significance (P-value *<*0.001). Please refer to Supplementary Data 1 for the complete list of the genes. Subsequently, we conducted gene set enrichment analyses (GSEA)^58^ utilizing the gene ontology (GO)^59^ and Kyoto Encyclopedia of Genes and Genomes (KEGG)^60^ datasets on the identified gene set. Notably, the GO terms related to lipoprotein particles emerged as significantly enriched within the gene set (Supplementary Fig. 2a). This observation is in line with prior studies that have implicated lipoprotein particles as significantly potential risk factors for Alzheimer’s diseasee^61–63^ and have highlighted the role of metabolic dysregulation in the progression of Alzheimer’s disease^64, 65^. Notably, the exploration of high-density lipoprotein-inspired treatments for Alzheimer’s disease has been a well-documented area of study^62, 63^. Fig. 4a illustrates the top 20 genes with the utmost statistical significance in the Mann-Whitney U test. Remarkably, 15 out of these 20 genes have been identified as potential risk factors for Alzheimer’s disease in previous studies. One notable example is *APOE*, which is the most prevalent highdensity lipoprotein in the central nervous system and has been consistently linked to Alzheimer’s disease in numerous studies^66–71^. Fig. 4b illustrates examples of the interactions from the GGI network between genes within the identified gene set. Please refer to Supplementary Data 2 for the complete list of the GGIs. Interestingly, aside from *APOE*, other genes within the *APOE* gene cluster, including *APOC1, APOC2*, and *APOC4*, were also identified as disease-related genes. This finding aligns with previous studies that have shown interdependent or independent associations of genes within the *APOE* gene cluster with Alzheimer’s disease^72–76^. For instance, it has been shown that the variant *APOE* and *APOC2* exhibit interactive effects on metabolic pathways, potentially contributing to the risk of Alzheimer’s disease^72^. *APOC1* also has been reported to serve as a risk factor for Alzheimer’s disease in combination with *APOE* ^74^. Furthermore, the combined effect of *APOE* and *CLU* on Alzheimer’s disease has been observed^77^. *SORL1* is an APOE receptor gene, which has been recognized as a genetic risk factor in Alzheimer’s disease. Recent research has elucidated the mechanistic connection between these two significant genetic factors in Alzheimer’s disease^78^. A neuron-specific interaction between Alzheimer’s disease risk factors *SORL1, APOE*, and *CLU* have also been shown in a recent study^79^. These observations highlight the proficiency of PRS-Net in not only identifying disease-related genes but also uncovering gene clusters that exhibit interactions contributing to the risk of Alzheimer’s disease.

**Fig. 2:**
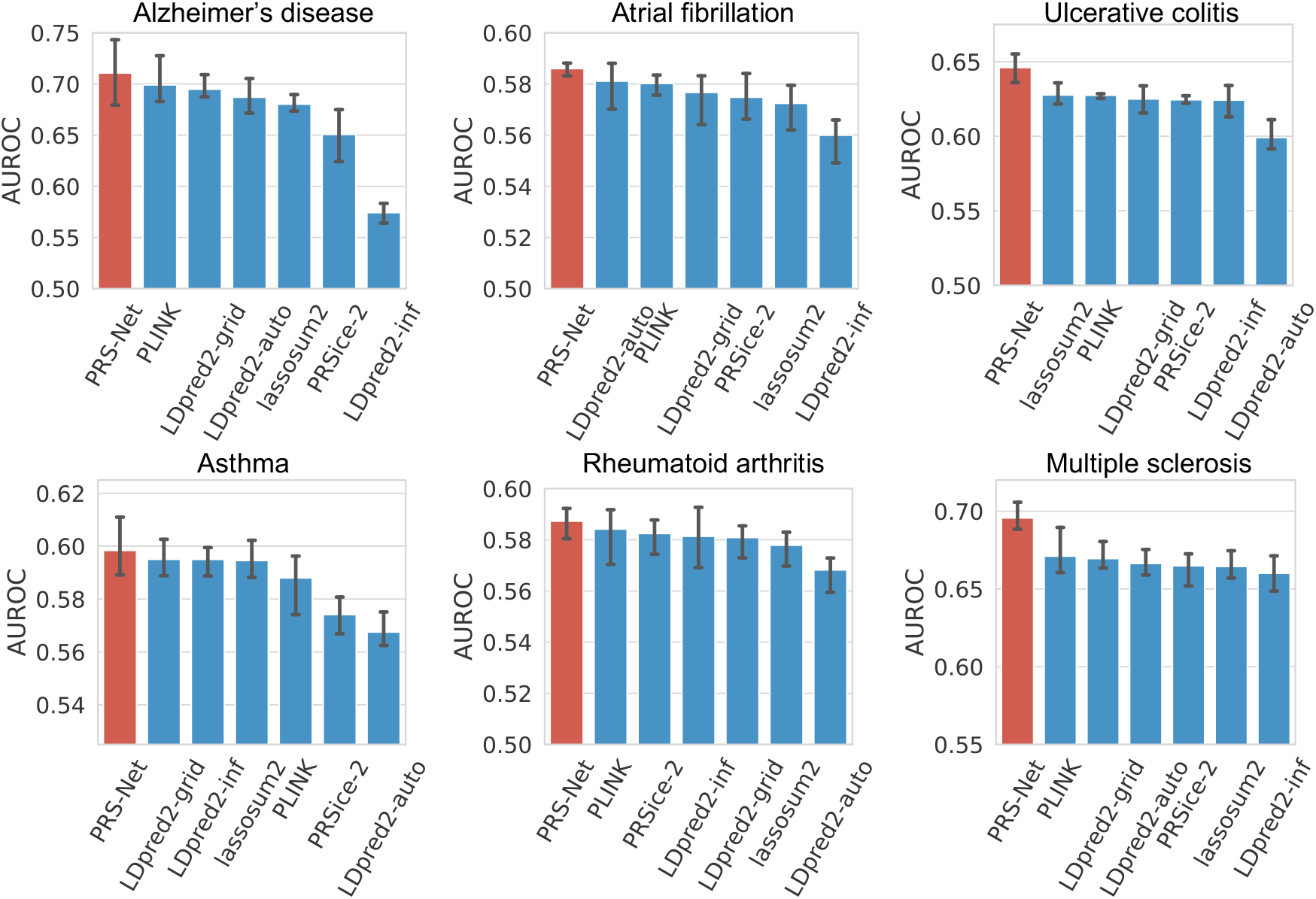
The PRS prediction performance of PRS-Net compared to baseline methods across a range of complex diseases, including Alzheimer’s disease, atrial fibrillation, ulcerative colitis, asthma, rheumatoid arthritis, and multiple sclerosis, measured in terms of the area under the receiver operating characteristic curve (AUROC). The bars are the estimated standard errors.

**Fig. 3:**
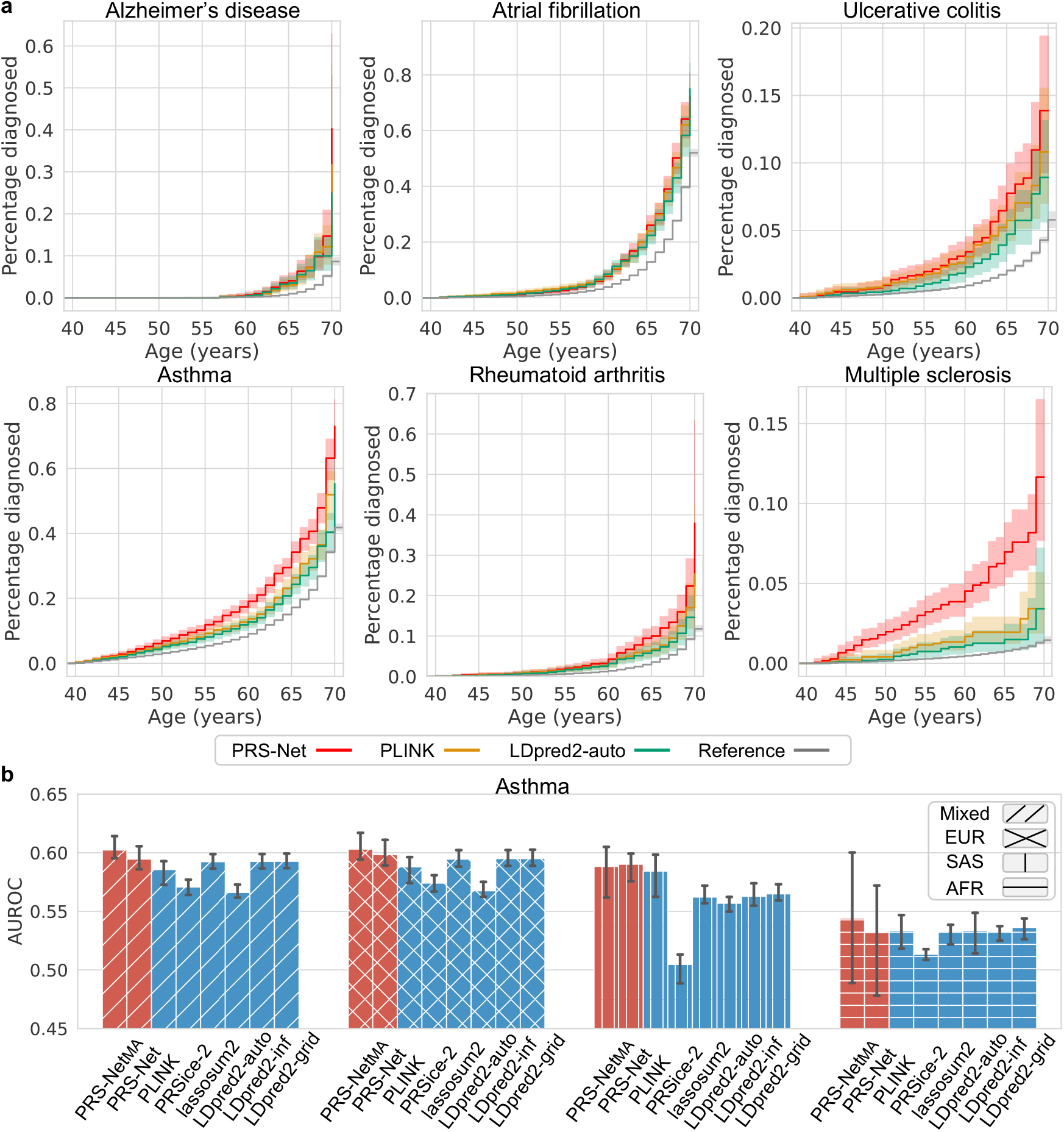
**a** The cumulative incidence plots of high-risk individuals (with the highest 5% PRSs) identified by PRS-Net and baseline methods. Each plot illustrates the estimated percentage of individuals diagnosed with a specific disease at different ages. We provide cumulative incidence plots for the original datasets as a reference. **b** The PRS prediction performance of PRS-Net compared to baseline methods on an asthma dataset encompassing multiple ancestries, including Western European (EUR), South Asian (SAS), and African (AFR) ancestry, measured in terms of the area under the receiver operating characteristic curve (AUROC). The results on the mixed ancestry test set are also reported. The bars are the estimated standard errors.

**Fig. 4:**
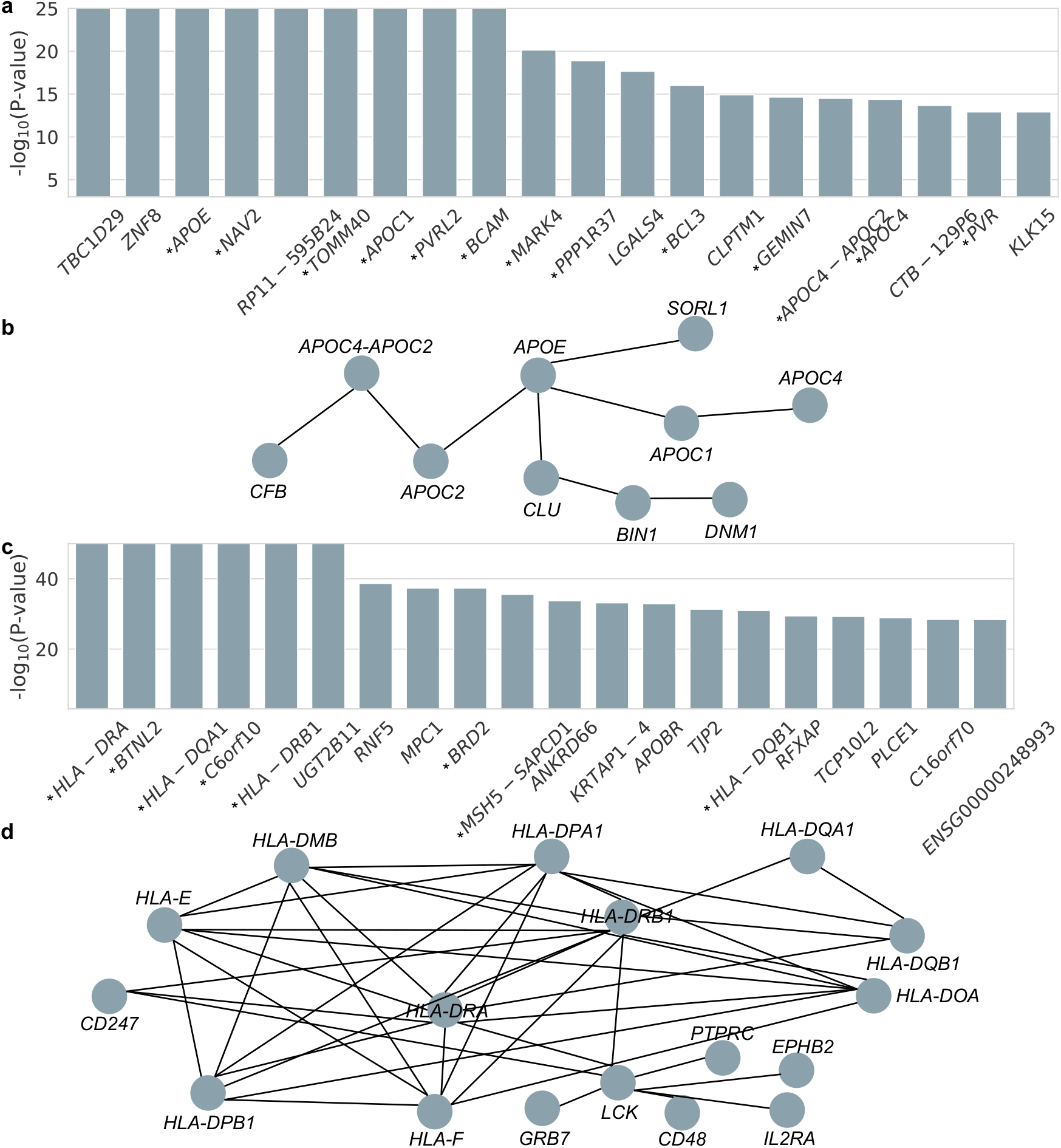
PRS-Net identifies disease-related genes and GGIs for Alzheimer’s disease and multiple sclerosis. **a** Top 20 genes with the highest statistical significance in the Mann-Whitney U test for Alzheimer’s disease. The Mann–Whitney U test was utilized to assess whether the attention scores for a particular gene among the cases were significantly higher than those observed in the control group. An asterisk preceding the gene name signifies that the gene has been reported to be associated with Alzheimer’s disease in previous studies. **b** Examples of interactions within the gene set with statistical significance (P-value *<*0.001) from the Mann-Whitney U test for Alzheimer’s disease. **c** Top 20 genes with the highest statistical significance in the Mann-Whitney U test for multiple sclerosis. **d** Examples of interactions within the gene set with statistical significance (Pvalue *<*0.001) from the Mann-Whitney U test for multiple sclerosis.

We also utilized PRS-Net to uncover genes and GGIs associated with multiple sclerosis. The Mann-Whitney U test identified a gene set with 456 potential risk genes (P-value *<*0.001). Please refer to Supplementary Data 3 for the complete list of the genes. The GSEA^58^ using the KEGG^60^ dataset on this gene set highlighted numerous immune-related pathways of statistical significance, such as antigen processing and presentation, allograft rejection, and graft-versus-host disease (Supplementary Fig. 4b). This finding aligns with the well-established understanding of multiple sclerosis as an autoimmune inflammatory disorder. The GSEA using the GO^59^ dataset, unveiled significant enrichment of GO terms related to the major histocompatibility complex (MHC) protein complex within the identified gene set (Supplementary Fig. 4a), which can be supported by previous studies that underscore the substantial genetic impact of MHCs on multiple sclerosis^80–84^. *HLA-DRA*, a subunit of *HLA-DR* which is a human MHC, was identified as the most significant gene in our analysis (Fig. 4c). Moreover, substantial *HLA* genes were identified as risk genes in our analysis (Fig. 4d). Please refer to Supplementary Data 4 for the complete list of the GGIs. This finding is in line with a previous study indicating that *HLA* interactions modulate genetic risk for multiple sclerosis^85^. Additionally, non-additive interactions between *HLA*s have been widely reported to significantly affect the risk of autoimmune diseases^38, 48–50^. These discoveries collectively provide compelling evidence of the potential of PRS-Net to offer valuable insights that advance our understanding of diseases.

### 3.3 Ablation studies

To assess the effectiveness of specific design choices in PRS-Net, we conducted comprehensive ablation studies. We introduced various modified frameworks derived from PRS-Net, each with distinct constraints: PRS-Net-GGI (omitting the GGI network), PRS-Net-Att+Sum (replacing the attentive readout module with a sum readout module, which summarized the node feature to derive the global-level representations), PRS-Net-Att+Mean (replacing the attentive readout module with a mean readout module, which computes the average of node features to derive global-level representations), and PRS-Net-Att+Max (replacing the attentive readout module with a max readout module, which extracts maximum values across node features to derive the global-level representations). We compared the performance of PRS-Net against these variants using datasets related to Alzheimer’s disease, multiple sclerosis, and ulcerative colitis. The results showcased that PRS-Net surpassed PRS-Net-GGI by an average relative improvement of 11.6%, underscoring the significance of incorporating the GGI network to capture the intricate genetic interactions associated with diseases (Fig. 5a). Furthermore, PRS-Net outperformed PRS-Net-Att+Sum, PRS-Net-Att+Mean, and PRS-Net-Att+Max with average relative improvements of 33.0%, 2.2%, and 8.4%, respectively, highlighting the effectiveness of the attentive readout module in summarizing node features (Fig. 5a).

**Fig. 5:**
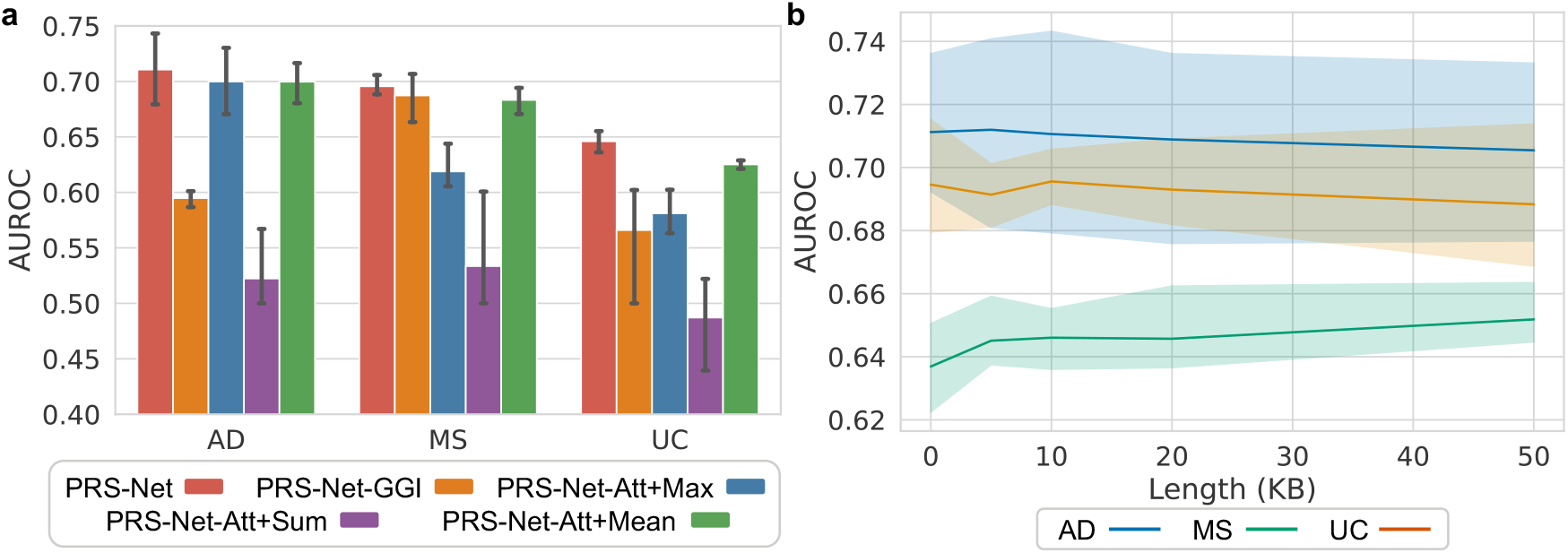
The results of ablation studies on PRS-Net. **a** The comparison results of PRS-Net and its variations, including PRS-Net-GGI (omitting the GGI network), PRS-Net-Att+Sum (replacing the attentive readout module with a sum readout module, which summarized the node feature to derive the global-level representations), PRS-Net-Att+Mean (replacing the attentive readout module with a mean readout module, which computes the average of node features to derive global-level representations), and PRS-Net-Att+Max (replacing the attentive readout module with a max readout module, which extracts maximum values across node features to derive the global-level representations), conducted on the datasets of Alzheimer’s disease (AD), multiple sclerosis (MS), and ulcerative colitis (UC). The bars are the estimated standard errors. **b** The PRS prediction performance of PRS-Net versus the extension lengths upstream and downstream of the transcription start and end sites.

Additionally, we explored the impact of varying extension lengths both upstream and downstream of the transcription start and end sites when calculating gene-level PRSs. We assessed different length values, including 0, 5, 10, 20, and 50 KB, and subsequently evaluated their prediction performance. The results demonstrated that PRS-Net is generally robust to different extension lengths (Fig. 5b). However, it is noteworthy that the performance of PRS-Net on the multiple sclerosis dataset significantly declined when the extension length was set to 0 KB (Fig. 5b). This observation suggested that including SNPs from non-coding regions can indeed enhance the accuracy of PRS prediction.

## Discussion

In this study, we develop PRS-Net, a deep-learning framework that offers interpretable and improved PRS predictions. By constructing a GGI network and incorporating a graph neural network, PRS-Net fully takes advantage of the power of non-linear associations between genetic factors and phenotypes. Additionally, the integration of an attentive readout module empowers PRS-Net to deliver interpretable predictions. Through comprehensive testing across six complex diseases, PRS-Net consistently achieved superior performance in comparison with baseline methods in PRS prediction. Furthermore, we demonstrated the interpretability of PRS-Net by using it to identify specific genes and GGIs that significantly impact the risk of Alzheimer’s disease and multiple sclerosis. In summary, PRS-Net provides a potent tool for accurate PRS prediction and biological discovery for complex diseases.

## Data availability

The GWAS data for Alzheimer’s disease can be accessed at https://ctg.cncr.nl/software/summarystatistics/. The GWAS data for atrial fibrillation can be accessed at https://cvd.hugeamp.org/downloads.html#summary/. The GWAS data for ulcerative colitis can be accessed at ftp://ftp.sanger.ac.uk/pub/project/humgen/summary_statistics/human/2016-11-07/. The GWAS data for asthma can be accessed at https://www.globalbiobankmeta.org/resources/. The GWAS data for rheumatoid arthritis can be accessed at https://data.cyverse.org/dav-anon/iplant/home/kazuyoshiishigaki/ra_gwas/ra_gwas-10-28-2021.tar/. The GWAS data for multiple sclerosis can be accessed at https://imsgc.net/?page_id=31/. The UKBB dataset is available at https://www.ukbiobank.ac.uk.

## Code availability

The source code of PRS-Net can be downloaded from the Github repository at https://github.com/lihan97/PRS-Net.

## Acknowledgments

This work was supported in part by the National Natural Science Foundation of China (T2125007 to J.Z.), the National Key Research and Development Program of China (2021YFF1201300 to J.Z.), the New Cornerstone Science Foundation through the XPLORER PRIZE (J.Z.), the Research Center for Industries of the Future (RCIF) at Westlake University (J.Z.), and the Westlake Education Foundation (J.Z.).

## Author contributions statement

H.L. and S.Z. conceived the concept and designed the study. H.L. and S.Z. developed the methodology and conducted data analysis. H.L., J.Z., M.S. and S.Z. are responsible for the data interpretation. S.Z., M.S. and J.Z. supervised the project. H.L. and S.Z. prepared the manuscript with the assistance from all other authors.

