## Supplementary material for "PRS-Net: Interpretable polygenic risk scores via geometric learning": SI.pdf

### 1 Supplementary notes

#### 1.1 Data quality control

**GWAS data quality control** In processing the GWAS data, several steps were taken. SNPs with a minor allele frequency (MAF) less than 0.1% and an imputation information score below 0.3 were excluded. In cases where SNPs had allelic inconsistencies between the base and target data, strand-flipping was applied if the inconsistency could be resolved; otherwise, the non-resolvable SNPs were removed. Duplicate SNPs were also eliminated to retain only one instance of each. Ambiguous SNPs were entirely removed from the dataset.

**Target data quality control** In processing the genotype-phenotype data, we implemented several filtering steps. We excluded SNPs with a genotyping rate below 1%, a minor allele frequency lower than 0.1%, or those not conforming to Hardy-Weinberg Equilibrium (with a P-value less than $1e^{-10}$). Duplicate SNPs were also eliminated to retain only one instance of each. Additionally, individuals with discrepancies between their reported sex and genetic sex were removed. We retained one individual from each pair sharing 3rd-degree kinship. For each phenotype, SNPs that exhibited a statistically significant difference between case and control genotype data (with a P-value less than $1e^{-5}$) were identified using Fisher's exact test based on case/control missing call counts for each variant and subsequently removed from the analysis.

#### 1.2 Training details

Our proposed method was implemented in PyTorch[1] version 1.13.1 and DGL[2] version 1.1.0 with CUDA version 11.6 and Python 3.7.16. We implemented a 1-layer GIN with a hidden size of 64. A 2-layer multi-layer perceptron is employed as a predictor. An AdamW optimizer with a learning rate $1e^{-4}$ was used to optimize the model. The training was performed with a batch size of 512 over a total of 20,000 steps, utilizing a single Nvidia A100 GPU.

#### 2 Supplementary Figures

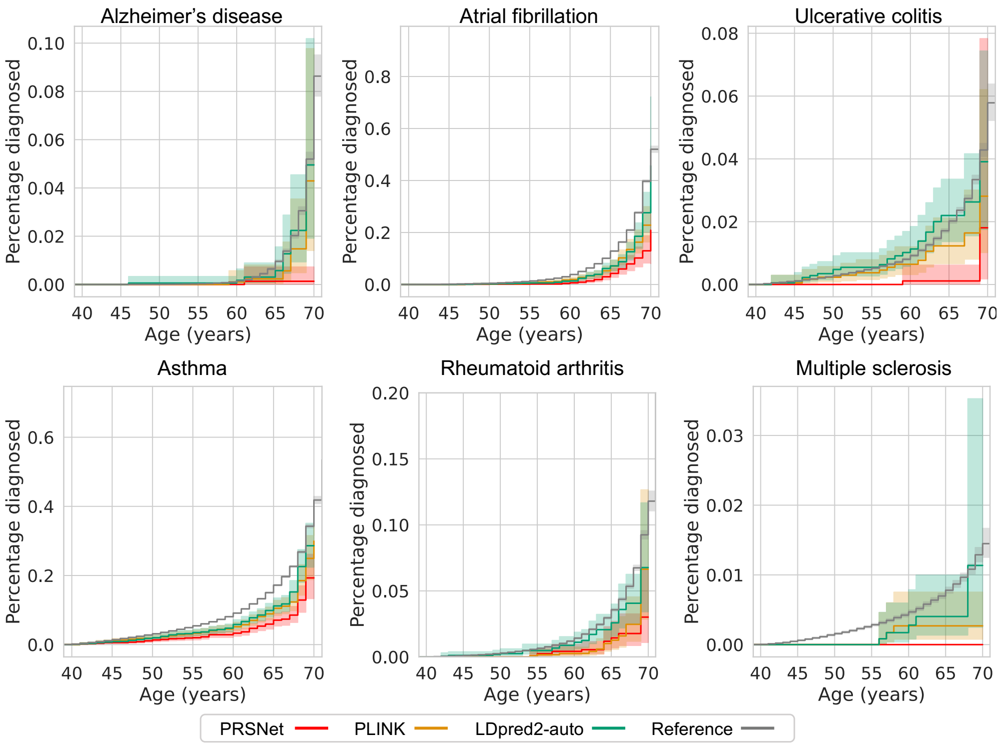


Supplementary Fig. 1: The cumulative incidence plots of low-risk individuals (with the lowest 5% PRSs) determined by PRS-Net and baseline methods. Each plot illustrates the estimated percentage of individuals diagnosed with a specific disease at different ages. We provide cumulative incidence plots for the original datasets as a reference.

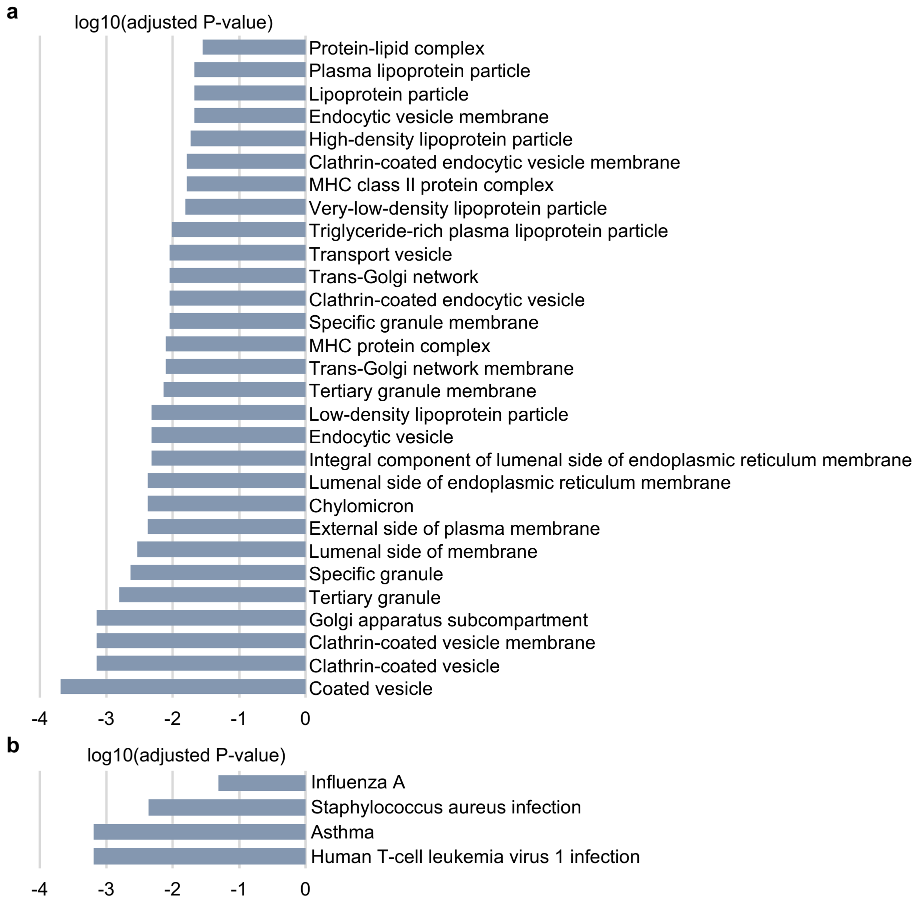


Supplementary Fig. 2: The results of gene set enrichment analysis (GSEA) for Alzheimer's disease, using (a) gene ontology (GO) and (b) Kyoto Encyclopedia of Genes and Genomes (KEGG) datasets, conducted on the gene set identified by the Mann–Whitney U test. The Mann–Whitney U test was employed to determine whether the attention scores of a specific gene for the cases significantly exceeded those of the control group.

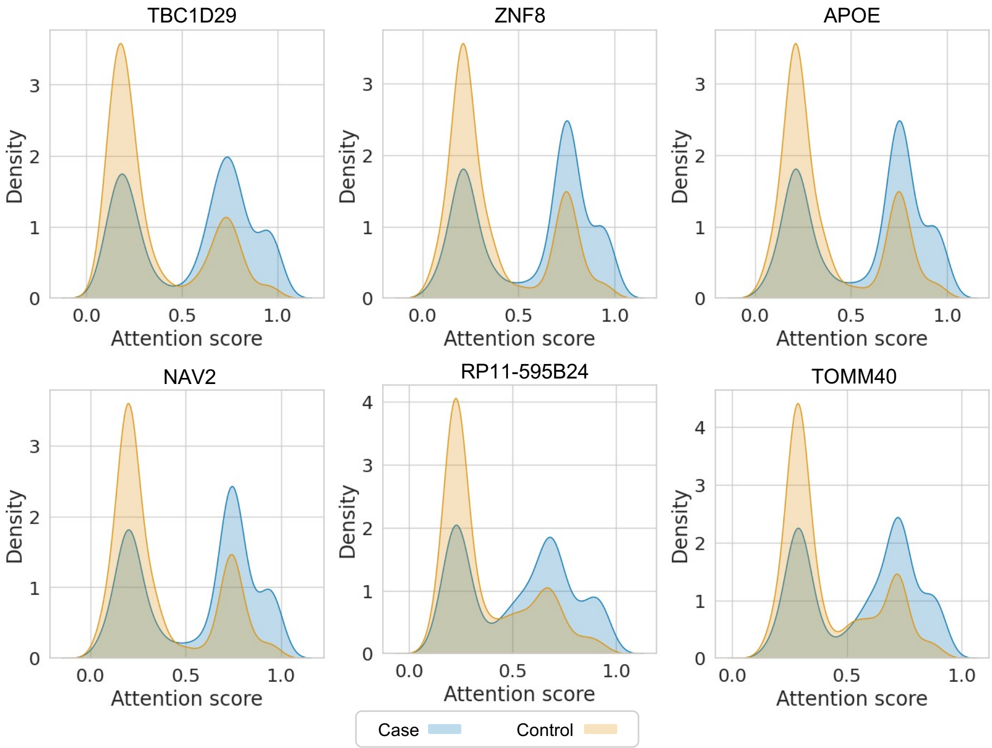


Supplementary Fig. 3: The kernel density estimate (KDE) plots illustrating the distributions of attention scores of the top 6 genes in the Mann–Whitney U test. The Mann–Whitney U test was employed to determine whether the attention scores of a specific gene for the cases significantly exceeded those of the control group.

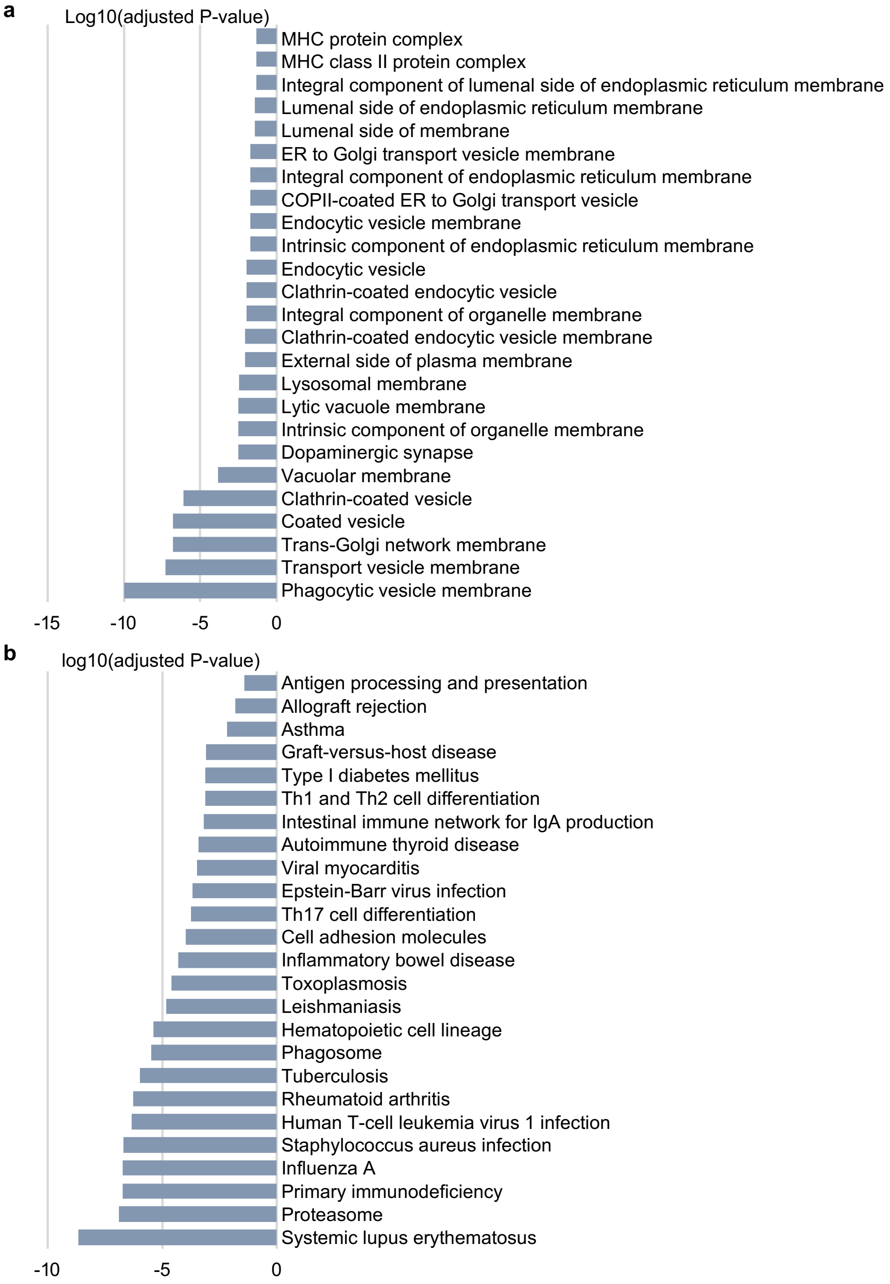


Supplementary Fig. 4: The results of gene set enrichment analysis (GSEA) for multiple sclerosis, using (a) gene ontology (GO) and (b) Kyoto Encyclopedia of Genes and Genomes (KEGG) datasets, conducted on the gene set identified by the Mann–Whitney U test. The Mann–Whitney U test was employed to determine whether the attention scores of a specific gene for the cases significantly exceeded those of the control group.

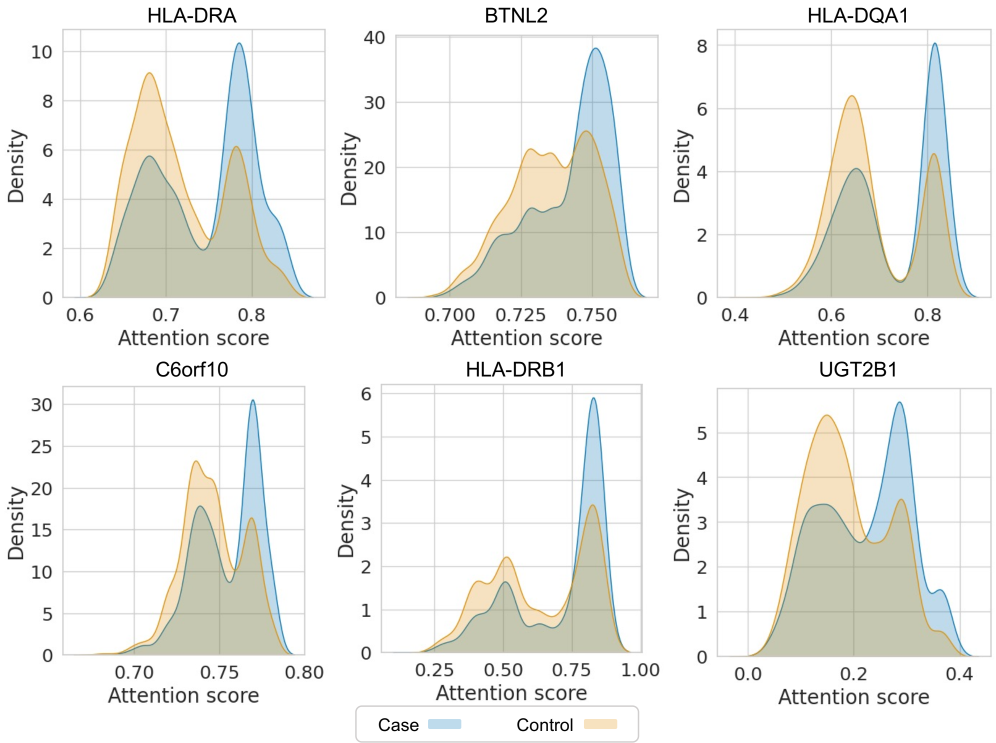


Supplementary Fig. 5: The kernel density estimate (KDE) plots illustrating the distributions of attention scores of the top 6 genes in the Mann–Whitney U test. The Mann–Whitney U test was employed to determine whether the attention scores of a specific gene for the cases significantly exceeded those of the control group.

#### 3 Supplementary Tables

Supplementary Table 1: ICD-10 codes used to identify different diseases.

| **Phenotype** | **ICD-10** |
|---|---|
| Alzheimer's disease | F00/G30 |
| Atrial fibrillation | I48 |
| Ulcerative colitis | K51/M07.5/M09.2 |
| Asthma | J45/J46 |
| Rheumatoid arthritis | M05/M06/M08.0 |
| Multiple sclerosis | G35 |

Supplementary Table 2: An overview of the distribution of various disesses across multiple population groups, including individuals of Western European (EUR), South Asian (SAS), and African (AFR) ancestry. Abbreviations: N (number), POS (positive), and NEG (negative).

| Phenotype | N.POS.EUR | N.NEG.EUR | N.POS.SAS | N.NEG.SAS | N.POS.AFR | N.NEG.AFR |
|---|---|---|---|---|---|---|
| Alzheimer's disease | 2310 | 388549 | 41 | 8772 | 53 | 8796 |
| Atrial fibrillation | 28754 | 362105 | 374 | 8439 | 266 | 8583 |
| Ulcerative colitis | 4293 | 386566 | 139 | 8674 | 51 | 8798 |
| Multiple sclerosis | 1742 | 389117 | 8 | 8805 | 16 | 8833 |
| Asthma | 41443 | 349416 | 1239 | 7574 | 1048 | 7801 |
| Rheumatoid arthritis | 7566 | 383293 | 242 | 8571 | 186 | 8663 |
